# A mathematical model of flagellar gene regulation and construction in *Salmonella enterica*

**DOI:** 10.1101/2020.01.29.924670

**Authors:** Kiersten Utsey, James P. Keener

## Abstract

Millions of people worldwide develop foodborne illnesses caused by *Salmonella enterica* (*S. enterica*) every year. The pathogenesis of *S. enterica* depends on flagella, which are appendages that the bacteria use to move through the environment. Interestingly, populations of genetically identical bacteria exhibit heterogeneity in the number of flagella. To understand this heterogeneity and the regulation of flagella quantity, we propose a mathematical model that connects the flagellar gene regulatory network to flagellar construction. A regulatory network involving more than 60 genes controls flagellar assembly. The most important member of the network is the master operon, *flhDC*, which encodes the FlhD_4_C_2_ protein. FlhD_4_C_2_ controls the construction of flagella by initiating the production of hook basal bodies (HBBs), protein structures that anchor the flagella to the bacterium. By connecting a model of FlhD_4_C_2_ regulation to a model of HBB construction, we investigate the roles of various feedback mechanisms. Analysis of our model suggests that a combination of regulatory mechanisms at the protein and transcriptional levels induce bistable FlhD_4_C_2_ levels and heterogeneous numbers of flagella. Also, the balance of regulatory mechanisms that become active following HBB construction is sufficient to provide a counting mechanism for controlling the total number of flagella produced.

**Author summary:** *Salmonella* causes foodborne illnesses in millions of people worldwide each year. Flagella, which are appendages that the bacteria use to move through the environment, are a key factor in the infection process. Populations of genetically identical bacteria have been observed to contain both motile cells, generally with 6–10 flagella, and nonmotile cells, with no flagella. In this paper, we use mathematical models of the gene network that regulates flagellar construction to explore how the bacteria controls the number of flagella produced. We suggest that a bacterium must accumulate a threshold amount of a master regulator protein to initiate flagella production and failure to reach the threshold results in no flagella. Downstream mechanisms that impact the amount of master regulator protein are sufficient to determine how many flagella are produced.

## Introduction

Millions of people worldwide develop foodborne illnesses caused by *Salmonella enterica* (*S. enterica*) every year [1]. The pathogenesis of *S. enterica* depends on flagella, which are appendages that the bacteria use to move through the environment. Flagella are located peritrichously on the bacterium’s surface and generally number between 6 and 10 per cell [2]. However, not all bacteria produce flagella. Partridge and Harshey describes the distributions of the number of flagella per bacterium under different growth conditions [3]. It was observed under each growth condition that some bacteria did not produce any flagella, while the number of flagella on bacteria that produced them roughly follows a normal distribution, where the mean depended on the growth condition. This phenotypic heterogeneity enables the evasion of the host immune system during acute infection [4]. It is not yet clear what mechanisms determine flagella heterogeneity and how the bacteria regulate the number of flagella produced.

Structurally, a flagellum is divided into three parts: basal body, hook, and filament. The basal body anchors the flagellum to the cell membrane, the hook is a flexible joint that links the rigid basal body with the filament, and the filament is a rigid helical structure which rotates to drive the cell forward. Assembly of functional flagella involves the temporally coordinated expression of more than 60 genes [5]. In particular, these genes are divided into three classes: Class I, Class II, and Class III depending on the timing of their activation. The *flhDC* operon is the only member of Class I. This operon encodes the master regulator proteins FlhD and FlhC, which form a transcriptional activation complex FlhD_4_C_2_ [6]. The FlhD_4_C_2_ complex is responsible for activation of Class II genes. Class II promoters control the expression of all the proteins required for the assembly of a functional hook-basal body (HBB) structure [7]. The HBB includes the flagellar type III secretion system, which exports flagellar proteins from the cytoplasm through the growing structure during assembly [8]. Once the HBB assembly is complete, FlhD_4_C_2_ activates expression of *flgM*, *fliA*, and other Class III genes. *fliA* encodes the flagella-specific sigma factor, *σ*^28^, which is essential for Class III promoter activation. *flgM* encodes an anti-sigma factor FlgM. Prior to HBB completion FlgM and *σ*^28^ form a complex that inhibits the activity of *σ*^28^. After the HBB is completed, the specificity of the secretion apparatus switches, so filament subunits and FlgM are secreted from the cell. *σ*^28^ facilitates the secretion of FlgM through the HBB, thereby acting as the FlgM type III secretion chaperone [9]. The secretion of FlgM frees *σ*^28^ inside the cell, which leads to the activation of Class III genes. Class III genes control the assembly of the filament, the chemotaxis system, and motor proteins.

While the components of the gene network have been identified, understanding the additional details and dynamics of the system requires further study. In particular, genetically identical bacteria grown in the same environment exhibit bistable expression of multiple flagellar genes [3]. Expression of Class II and Class III genes, *flhB* and *fliC*, respectively, is bimodal, and this bimodality is likely due to bistability upstream in the network [10–13]. Other factors also impact the flagella gene network, including nutrient availability, which enhances the expression of flagellar and motility genes [14]. Nutrients repress RflP, formerly YdiV, expression through the protein CsrA [15]. Low nutrient levels correspond to high RflP expression, and high nutrient levels correspond to low RflP expression. The tuning of RflP expression influences the flagellar gene network because RflP promotes the degradation of FlhD_4_C_2_ through the protease ClpXP [16].

A series of mathematical models have been proposed to explore the dynamics of the flagellar gene network [17–19]. While these models investigated the roles of some of the feedback mechanisms in the network, they fail to describe the heterogeneity observed in bacterial populations, since all bacteria are predicted to grow flagella. Further, the number of HBBs was modeled as a continuous variable, although HBBs are discrete entities.

Meanwhile, Koirala et al. presented a model that attempted to explain how nutrients tune flagellar gene expression dynamics in *Salmonella enterica* [13]. Using experimental observations and a deterministic model, the authors demonstrated that environmental nutrient levels can regulate RflP expression and that this expression can lead to bistable expression of Class II and Class III genes in the bacteria [13]. The model does not include HBB construction, or, consequently, the additional feedback mechanisms that are activated following HBB completion.

To investigate the heterogeneous flagella quantities within populations of clonal bacteria, we propose two mathematical models of the flagellar gene regulatory network. We first consider a sub-system of the network to focus on FlhD_4_C_2_ production and degradation. We use bifurcation analysis of an ordinary differential equation model to explore the role of *flhDC* and *rflP* expression rates on overall FlhD_4_C_2_ concentration. We find bistability in FlhD_4_C_2_ concentration as a function of both expression rates. We then extend the model to consider HBB construction and additional regulatory mechanisms. We retain the bistability observed in the sub-system model and investigate factors that impact the bistable regime. We also propose a mechanism for how *S. enterica* controls the number of flagella produced via FliT:FliD dynamics and Class III regulation of FliZ.

## Models

### Sub-system model

We first consider a model of the key components that regulate *flhDC* expression and FlhD_4_C_2_ protein concentrations. The model includes the FliZ-RflP-FlhD_4_C_2_ feedback loop and RflM activity. Expression of *fliZ* is up-regulated by FlhD_4_C_2_ and FliZ represses expression of *rflP* [20]. RflP inhibits the activity of FlhD_4_C_2_ by binding to the FlhD subunit and targeting FlhD_4_C_2_ for proteolysis via ClpXP [16]. Thus, FliZ is involved in a negative-negative feedback loop through RflP on the FlhD_4_C_2_ protein. FlhD_4_C_2_ up-regulates expression of *rflM*, and RflM represses the transcription of FlhD_4_C_2_ [21–23]. Figure 1 summarizes the mechanisms included in this model.

**Fig 1.**
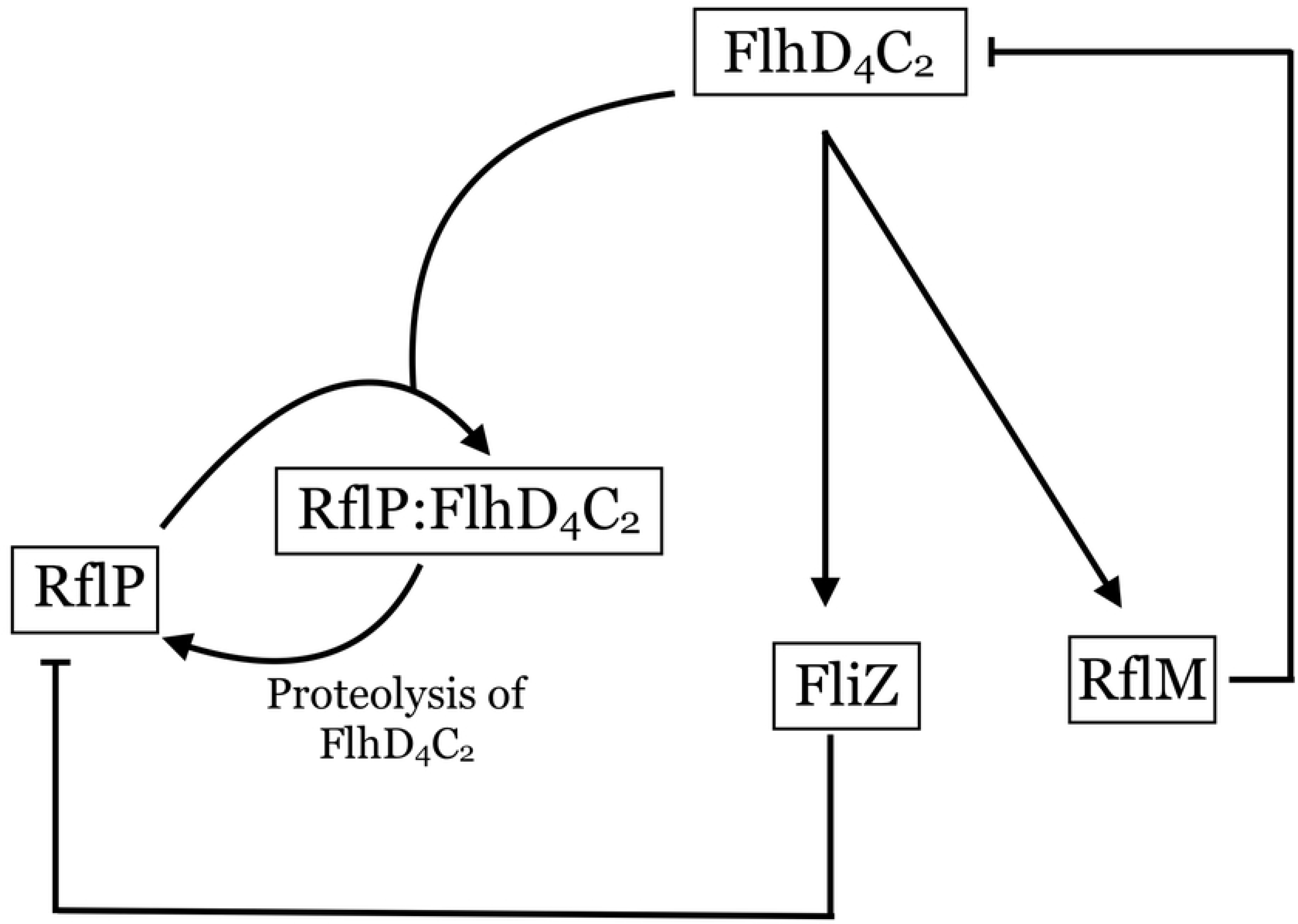
Sub-system of gene network. FlhD_4_C_2_ upregulates FliZ and RflM. The formation of the RflP:FlhD_4_C_2_ complex leads to FlhD_4_C_2_ proteolysis by ClpXP and RflP recycling. The production of RflM leads to repression of *flhDC* expression.

We set *x* =[FlhD_4_C_2_], *z* =[FliZ], *y* =[RflP], *c*_1_ =[RflP:FlhD_4_C_2_] and *m* =[RflM], and we construct the following model

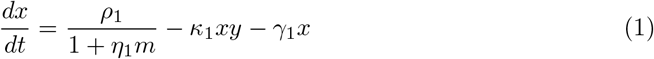

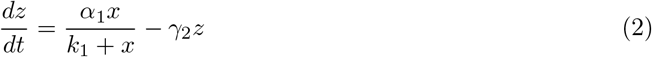

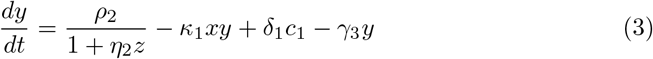

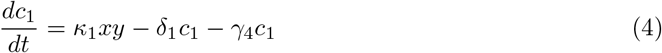

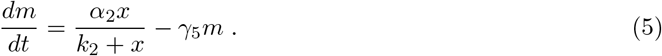

We assume the system is well-mixed, and we model transcription and translation processes as Michaelis-Menten functions and protein binding, unbinding, and degradation processes linearly. FlhD_4_C_2_ is produced at a basal rate of *ρ*_1_, and the production rate is divided by the concentration of RflM, scaled by *η*_1_. FlhD_4_C_2_ binds to RflP at rate *κ*_1_. FliZ is produced as a function of FlhD_4_C_2_ concentration. RflP is produced at a basal rate of *ρ*_2_, and the production is repressed by FliZ. External nutrient availability also represses *rflP* expression, so *ρ*_2_ is a proxy for nutrient availability [15]. RflP forms a complex with FlhD_4_C_2_ and is recycled at rate *δ*_1_ where FlhD_4_C_2_ is degraded by ClpXP. We assume that the complex does not dissociate spontaneously. RflM is produced as a function of FlhD_4_C_2_ concentration. All components, including the RflP:FlhD_4_C_2_ complex, are assumed to degrade at a small, basal rate, *γ*_*i*_, where *i* = 1, 2, …, 5. Table 1 lists the model parameters with descriptions and values.

**Table 1.** Sub-system model parameters.

| Parameter | Description | Value |
| --- | --- | --- |
| $\rho_1$ | Production rate of FlhD<sub>4</sub>C<sub>2</sub> | 0–5 |
| $\rho_2$ | Maximum production rate of RflP | 0–2 |
| $\alpha_1$ | Maximum production rate of FliZ | 2 |
| $\alpha_2$ | Maximum production rate of RflM | 1 |
| $k_1$ | Michaelis constant for FliZ expression | 5 |
| $k_2$ | Michaelis constant for RflM expression | 1 |
| $\kappa_1$ | Rate of binding of RflP and FlhD<sub>4</sub>C<sub>2</sub> | 4 |
| $\eta_1$ | Repression coefficient of RflM on FlhD<sub>4</sub>C<sub>2</sub> expression | 0.01 |
| $\eta_2$ | Repression coefficient of FliZ on RflP expression | 0.5 |
| $\gamma_1$ | Degradation rate of FlhD<sub>4</sub>C<sub>2</sub> | 0.1 |
| $\gamma_2$ | Degradation rate of FliZ | 0.1 |
| $\gamma_3$ | Degradation rate of RflP | 0.1 |
| $\gamma_4$ | Degradation rate of RflP:FlhD<sub>4</sub>C<sub>2</sub> complex | 0.01 |
| $\gamma_5$ | Degradation rate of RflM | 0.1 |
| $\delta_1$ | ClpXP degradation rate of FlhD<sub>4</sub>C<sub>2</sub> | 0.1 |

Parameter values were chosen to capture the desired dynamics of the system. Concentration and time both have arbitrary units.

#### Bifurcation analysis of sub-system model

Since some bacteria produce flagella and others do not, we suspect that FlhD_4_C_2_ concentration is bistable. We use resultant analysis of the model to determine what parameter regimes support bistability [24]. To apply resultant analysis, we take the model to steady state, reduce the system to a single polynomial in one variable, and perform a rescaling to reduce the number of parameters. We then solve the equation for a rescaled parameter of interest. To find double roots of the first derivative, we then take the resultant of the first and second derivatives of the right side of the equation. The curve in parameter space on which the first and second derivatives have a common root corresponds to the separation between monotonic and triphasic behavior. We first take the model to steady state and find

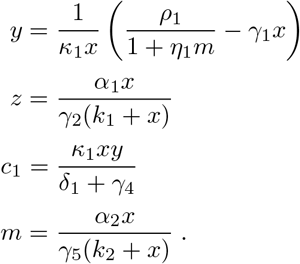

Substituting into 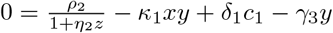, we find

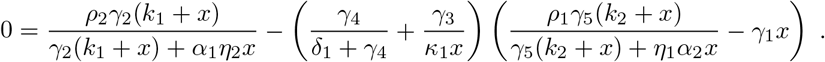

We reduce the number of parameters by taking the rescaling *x* = *k*_1_*w* and letting 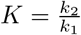, 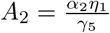, 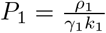, 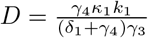, 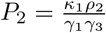 and 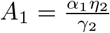. We find

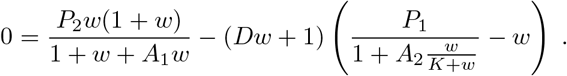

We solve for *P*_2_ to find

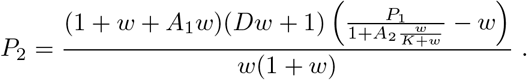

We define *f*(*w*) as

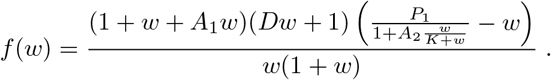

We next use resultant analysis of *f*′(*w*) and *f*″(*w*) to determine when *f*(*w*) is monotone and when it is triphasic. The triphasic regions correspond to bistability, and the monotone regions correspond to monostability.

We first consider the case in which *η*_1_ = 0 (*A*_2_ = 0), which means that RlfM does not impact *flhDC* expression. *f*(*w*) simplifies to

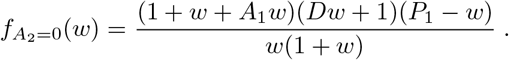

We next calculate the resultant of the numerators of the first and second derivatives of 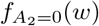 using Maple (2019; Maplesoft, Waterloo, ON), and we find

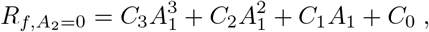

where

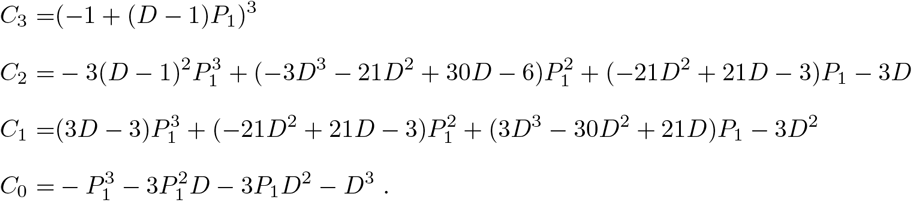

The function 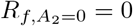 separates parameter space into regions for which 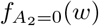 is monotone increasing (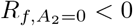) and triphasic (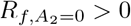). In the triphasic region, there is bistability, which will be verified later. The solid curve in Fig 2 shows the zero level contour for 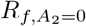.

**Fig 2.**
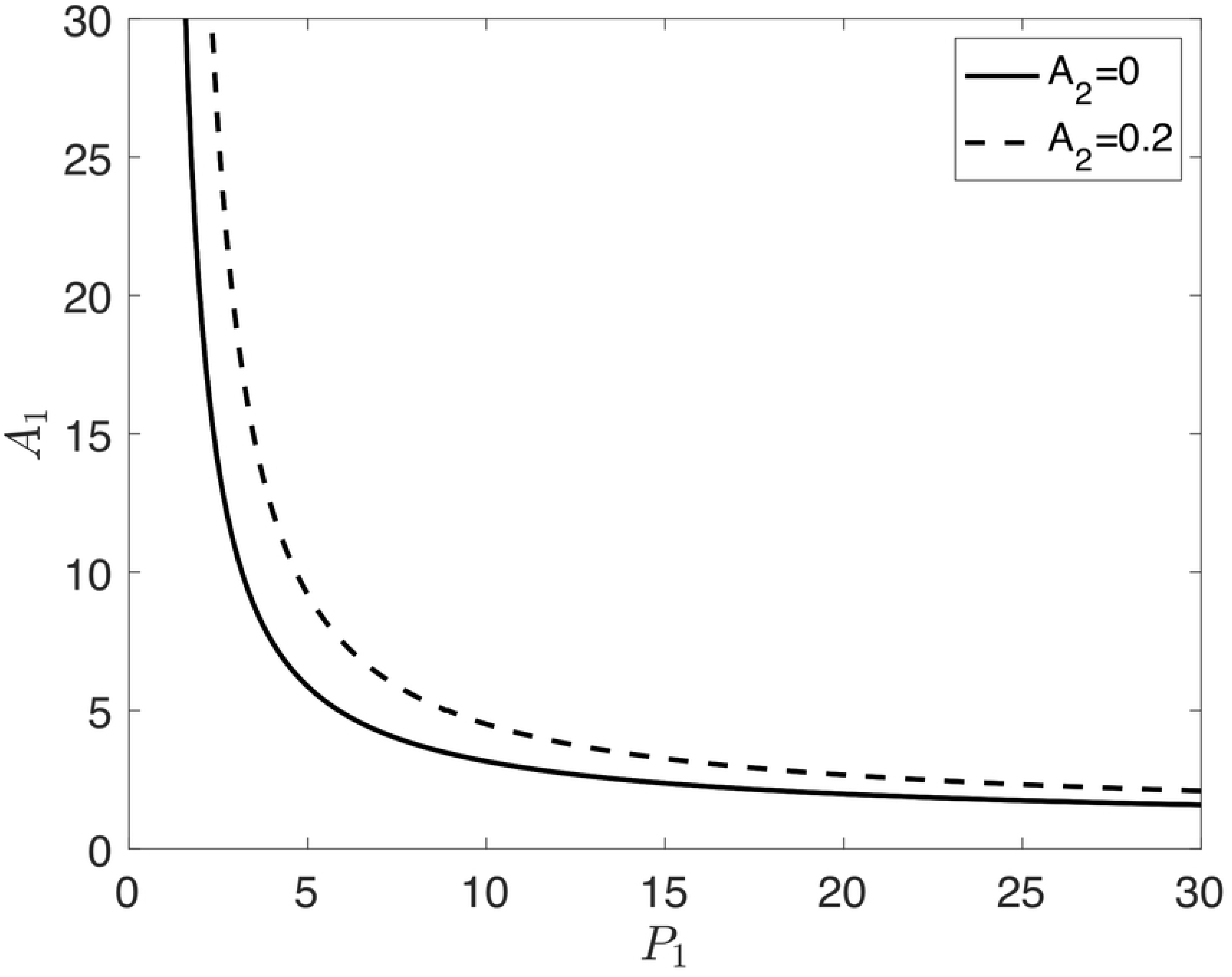
Zero level contours of *R*_*f*_, where *D* = 4 and *K* = 0.2. The solid curve indicates the *A*_2_ = 0 case, and the dashed curve indicates the *A*_2_ = 0.2 case. *R*_*f*_ < 0 below and to the left of the curve and *R*_*f*_ > 0 above and to the right of the curve.

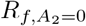 is negative to the left of the curve and is positive to the right of the curve. *f*(*w*) is monotone for values of *P*_1_ and *A*_1_ that fall to the left of the curve, whereas for values that fall to the right of the curve, *f*(*w*) is triphasic, and there is bistability. Thus, bistability is possible under constitutive expression of *flhDC* (i.e. when *A*_2_ = 0).

We note that in the case that *D* = 0, the resultant reduces to

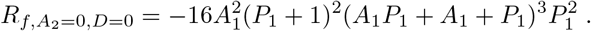

For positive values of *A*_1_ and *P*_1_, 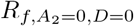 is always negative, and either *A*_1_ = 0 or *P*_1_ = 0 satisfy *R*_*f*_ = 0. This indicates that we must have *D* > 0 to have bistability. In terms of the original parameters, this means that *γ*_4_ > 0 is required to have bistability.

We next consider the case in which *η*_1_ ≠ 0 (*A*_2_ ≠ 0), meaning RflM represses the expression of *flhDC*. We calculate the resultant of the numerators of the first and second derivatives of *f*(*w*), but it is too long to show here. The dashed curve in Fig 2 is the zero level contour for *R*_*f*_ when RflM activity is included. We see that the *R*_*f*_ = 0 curve moves up and to the right as *A*_2_ increases, which decreases the size of the parameter space in which there is bistability. In particular, the repressive activity of RflM requires an increase in the baseline production rates of FlhD_4_C_2_ (*P*_1_) and FliZ (*A*_1_) to have bistability.

We next use the parameters in Table 1 and XPPAUT (8.0) to visualize the bistable regime identified by the resultant analysis. Fig 3 shows the steady state concentration of FlhD_4_C_2_ plotted as a function of the parameters *ρ*_1_ and *ρ*_2_, demonstrating bistability for certain regions of *ρ*_1_, *ρ*_2_ parameters. Fig 3 also shows that the value of the FlhD_4_C_2_ production term, 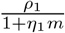, is multivalued for the same regions of *ρ*_1_ and *ρ*_2_.

**Fig 3.**
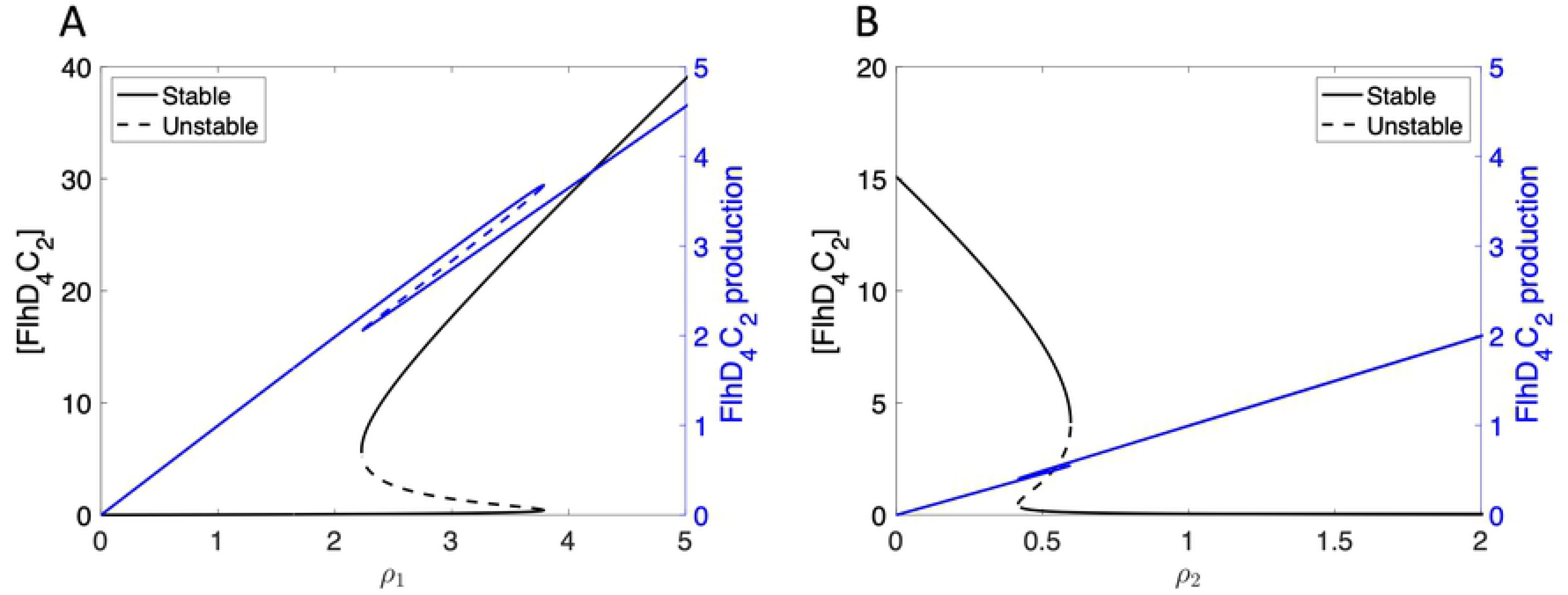
Steady state solution for concentration of FlhD_4_C_2_ (black) and FlhD_4_C_2_ production rate (blue) plotted as functions of *ρ*_1_ and *ρ*_2_. (A) Bistability is exhibited as *ρ*_1_ varies, where *ρ*_2_=1. (B) Bistability is also found as *ρ*_2_ varies and *ρ*_1_=1.5 is fixed.

Depending on the values of *ρ*_1_ and *ρ*_2_, the steady state value of FlhD_4_C_2_ concentration has one solution or three solutions. As shown in Fig 3A, the FlhD_4_C_2_ concentration is low for small *ρ*_1_ values. This means that the bacteria do not produce enough FlhD_4_C_2_ to initiate flagella production. With *ρ*_1_ increased sufficiently, a saddle node bifurcation occurs, and bistability arises. The lower solution represents the case in which the bacterium produces no flagella, and the upper solution represents the case in which the bacterium produces flagella. Increasing *ρ*_1_ further results in another saddle node bifurcation, and a return to a single steady state value for FlhD_4_C_2_ concentration. The expectation is that for high *flhDC* production rates, the bacterium will have a high FlhD_4_C_2_ concentration, and, consequently, produce flagella. Saddle node bifurcations are also observed for *ρ*_2_, as shown in Fig 3B. In this case, small *ρ*_2_ values result in a high steady state FlhD_4_C_2_ concentration, intermediate *ρ*_2_ values result in bistability, and large *ρ*_2_ values result in a low FlhD_4_C_2_ concentration.

Many factors control *flhDC* expression, so it is reasonable to expect that clonal bacteria may have different values for *ρ*_1_. In this case, the variation in *ρ*_1_ values could contribute to heterogeneity in flagella number, assuming the bacteria on the lower branch have no flagella and bacteria on the upper branch have enough FlhD_4_C_2_ to produce flagella. Also, since nutrient availability represses *rflP* expression, small variations in nutrient availability could result in differing values for *ρ*_2_. Variability in *ρ*_2_ would result in different concentrations of FlhD_4_C_2_, and, consequently, the presence or absence of flagella. We note that, as predicted by the resultant analysis, *γ*_4_ > 0 is required for bistability (not shown).

We also consider the *η*_1_ = 0 case, which corresponds to a knockout of RflM or constitutive expression of *flhDC* via an inducible promoter. Under this condition, the FlhD_4_C_2_ production term reduces to *ρ*_1_. Fig 4 shows the steady state concentration of FlhD_4_C_2_ is bistable for certain regions of *ρ*_1_ and *ρ*_2_, but the FlhD_4_C_2_ production term is not multivalued.

**Fig 4.**
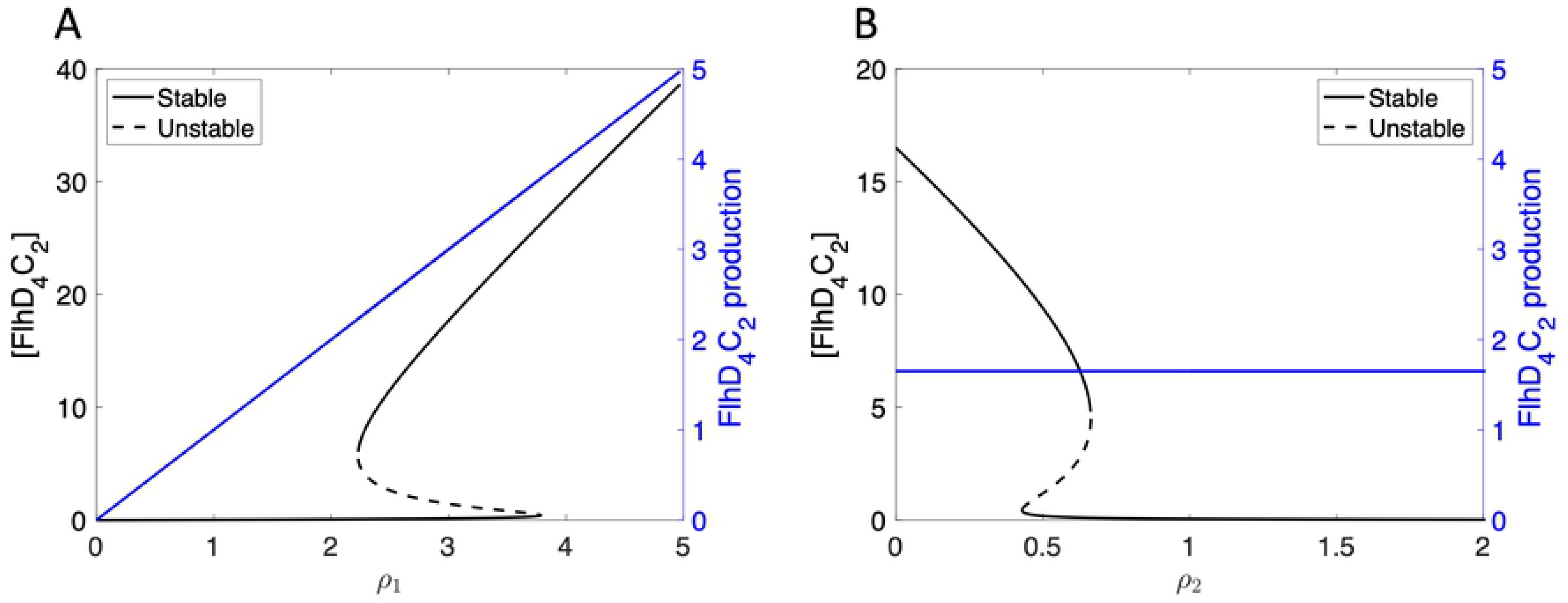
Steady state solution for concentration of FlhD_4_C_2_ (black) and FlhD_4_C_2_ production rate (blue), where *η*_1_=0. Bistability is exhibited for FlhD_4_C_2_ concentration, but not FlhD_4_C_2_ production, as *ρ*_1_ varies, where *ρ*_2_=1 (A), and as *ρ*_2_ varies and *ρ*_1_=1.65 (B).

In this case, constitutive expression of *flhDC* can lead to bistable FlhD_4_C_2_ levels. In particular, the *η*_1_ = 0 case shows bistability in FlhD_4_C_2_ even when the production of *flhDC* is not multivalued. Therefore, we suggest that RflM knockout cells should still exhibit bistability in the protein level of FlhD_4_C_2_.

In anticipation of a need to understand the role of FlhD_4_C_2_ degradation, we next investigate the impact of the basal degradation rate of FlhD_4_C_2_ on FlhD_4_C_2_ concentration. Fig 5A shows bistability in FlhD_4_C_2_ concentration as a function of the parameter *γ*_1_.

**Fig 5.**
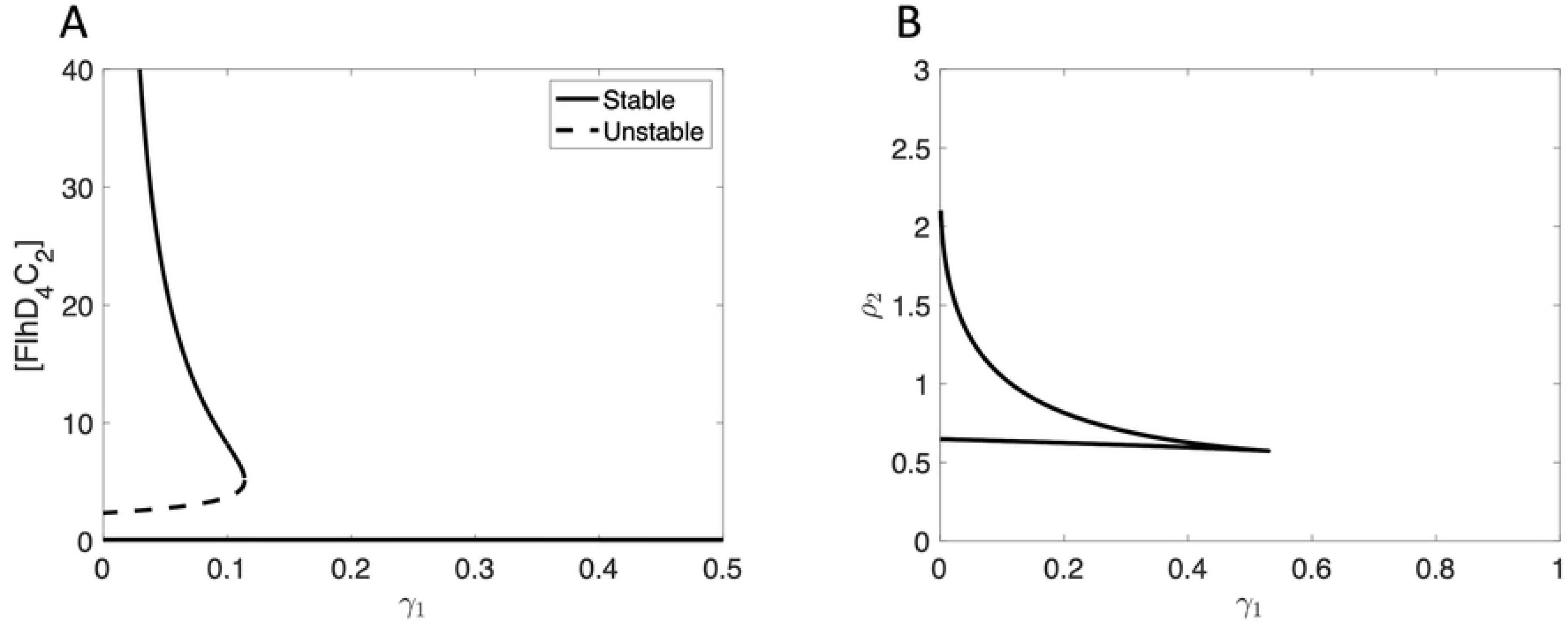
Steady state solution and two parameter bifurcation diagram, where *ρ*_1_ = 2.5. (A) Steady state solution for concentration of FlhD_4_C_2_, where *ρ*_2_ = 1. Bistability is seen as *γ*_1_ varies. (B) Two parameter bifurcation diagram. The region of parameters within the curve support bistable FlhD_4_C_2_ steady state concentrations, whereas parameter values above and below the curve result in a single steady state for FlhD_4_C_2_ concentration.

A saddle node bifurcation is observed for FlhD_4_C_2_ concentration as a function of *γ*_1_. Small *γ*_1_ values result in bistable steady state FlhD_4_C_2_ concentration, and increasing *γ*_1_ sufficiently results in a low FlhD_4_C_2_ concentration. We next investigate the impact of *γ*_1_ on the bistability observed in FlhD_4_C_2_ as a function of *ρ*_2_. Fig 5B shows a cusp bifurcation in the *γ*_1_-*ρ*_2_ plane. This demonstrates that the bistability previously observed in FlhD_4_C_2_ as a function of *ρ*_2_ is sensitive to *γ*_1_.

### Extended model

To explore how the bacteria regulate the total number of flagella constructed, we extend the model to consider the construction of HBBs and additional regulatory mechanisms that become active following HBB completion. In particular, FlhD_4_C_2_ up-regulates the *fliDST* operon, which encodes the FliT and FliD proteins. Before HBB completion, FliT and FliD are bound together, which inhibits the activity of FliT. Following HBB completion, FliD is secreted through the HBB and is assembled as the cap of the flagellum. The secretion of FliD frees FliT in the cell to bind to FlhD_4_C_2_ and target FlhD_4_C_2_ for proteolysis via ClpXP [25–27]. FliT is recycled during FlhD_4_C_2_ degradation [25]. FlhD_4_C_2_ also up-regulates production of FlgM and *σ*^28^, which form a complex, and complete HBBs secrete FlgM, resulting in free *σ*^28^ in the bacterium. *σ*^28^ up-regulates the expression of Class III genes, including *fliC*. FliC is also secreted through the HBB to become the flagellar filament. *σ*^28^ also up-regulates *fliZ*, which means *fliZ* is under both Class II and Class III regulation. The gene network for the full model is shown in Fig 6.

**Fig 6.**
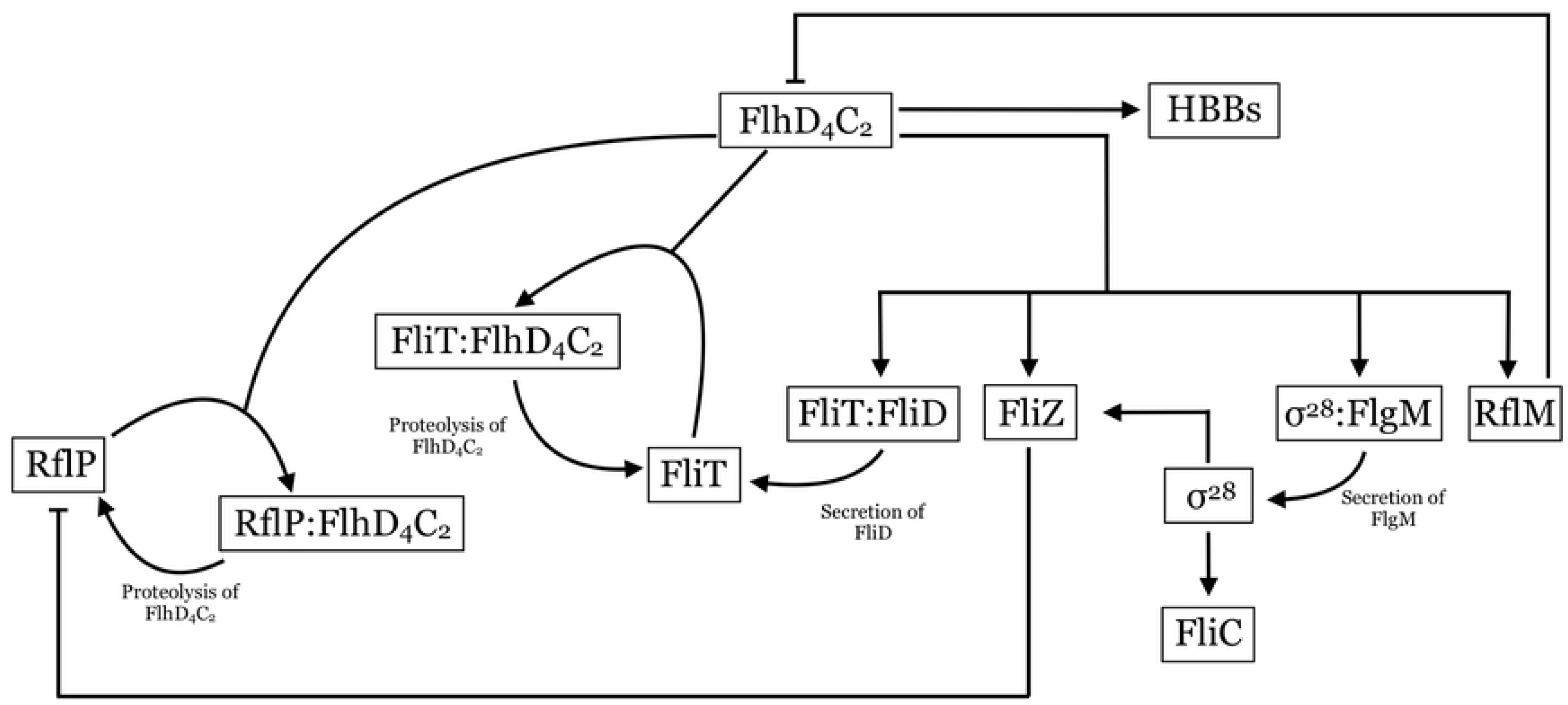
Full gene network. Arrows indicate positive regulation or protein binding/unbinding processes. Blunt arrows indicate repression of gene expression.

For simplicity, we assume that FliT and FliD are produced as a complex (FliT:FliD), rather than modeling the production of each protein and formation of the FliT:FliD complex. Similarly, *σ*^28^ and FlgM are assumed to be produced as the complex *σ*^28^:FlgM.

We use the variables defined in the sub-system model and set *v* =[FliT], *c*_2_ =[FliT:FlhD_4_C_2_], *c*_3_ =[FliT:FliD], *c*_4_ =[*σ*^28^:FlgM], *a* =[*σ*^28^], and *f* =[FliC]. Referring to Fig 6, we construct the following model

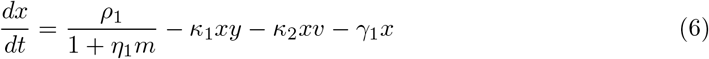

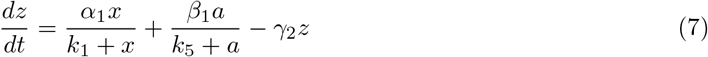

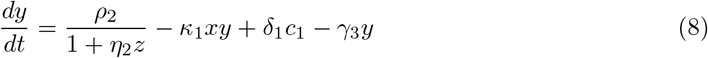

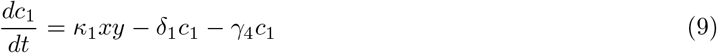

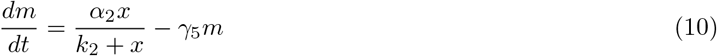

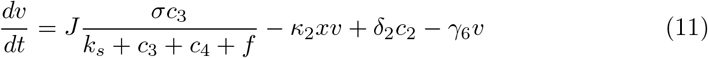

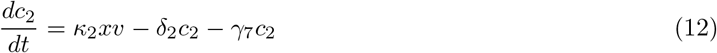

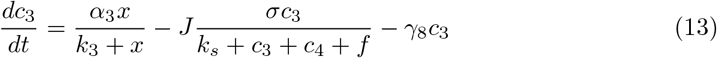

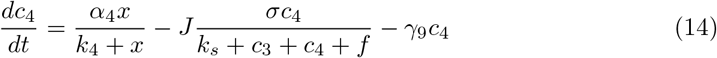

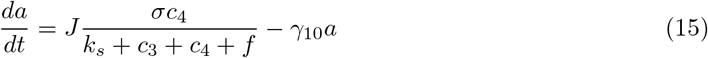

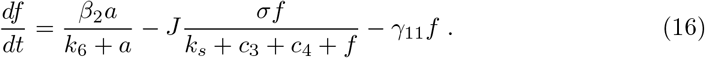

As before, we assume that the system is well-mixed, and we model transcription and translation processes as Michaelis-Menten functions and protein binding, unbinding, and degradation processes linearly. *J* is the number of complete HBBs that are capable of secreting FlgM, FliD, and FliC, and we assume that the secretion of the three components, FlgM, FliD, and FliC, through the HBBs is competitive and saturating. We assume that these three components have the same secretion affinity, *σ*. As in the sub-system model, we assume that every compound, including the complexes, degrade at some basal rate.

#### Model reduction

The model involves both protein level dynamics, including binding and degradation, and transcription and translation. We assume that the protein level dynamics occur on a faster time scale than transcription and translation, so we take a quasi-steady-state approximation for the protein level dynamics. We assume that *c*_1_ =[RflP:FlhD_4_C_2_], *y* =[RflP], and *c*_2_ =[FliT:FlhD_4_C_2_] are fast equilibrating, so that they are at equilibrium. We therefore assume that

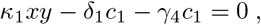

meaning that

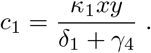

We also assume that

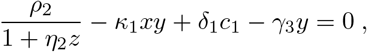

meaning that

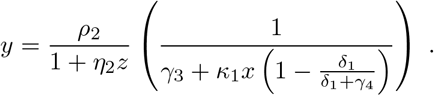

We now let *u* = *v* + *c*_2_, the sum of [FliT] and [FliT:FlhD_4_C_2_], so we have

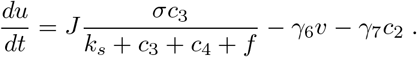

We then assume that

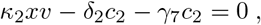

meaning that

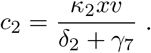

Then we have

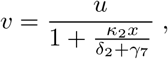

so that

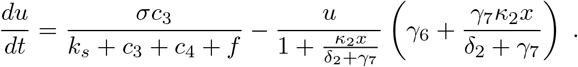

Using the quasi-steady-state approximation, the model no longer includes equations for *c*_1_ =[RflP:FlhD_4_C_2_], *y* =[RflP], and *c*_2_ =[FliT:FlhD_4_C_2_]. The reduced model with competitive, saturating secretion, is

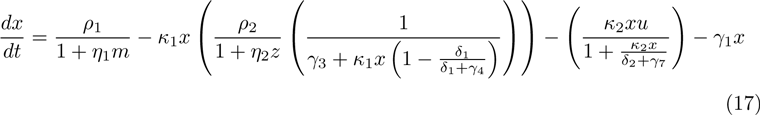

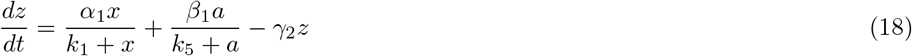

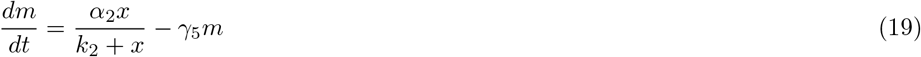

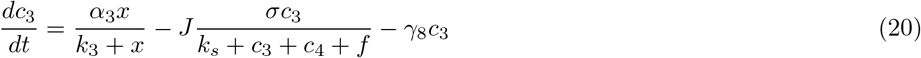

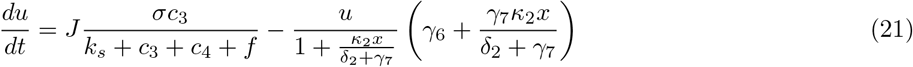

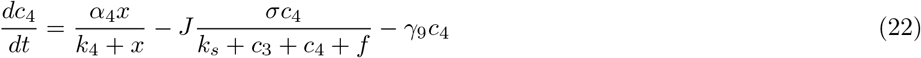

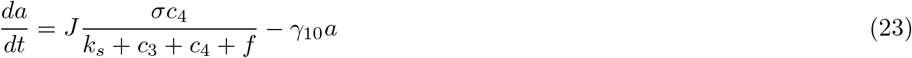

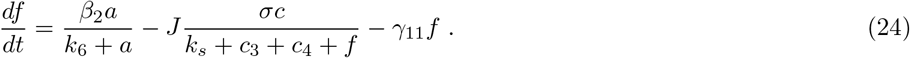

#### HBB model

Since little is understood about HBB initiation and construction, we propose a simple model that assumes the environment within the bacteria is well-mixed. We assume that a general HBB protein is produced at a rate that depends on the concentration of FlhD_4_C_2_ in the bacterium. Mouslim and Hughes showed that flagella construction requires a threshold of *flhDC* expression, and, consequently, Class II expression [21]. This finding suggests that the concentration of HBB proteins must reach some threshold before initiation of HBB construction. We assume that once the concentration of the HBB protein reaches a threshold, nucleation of a HBB occurs, and the free HBB protein concentration drops to zero. The newly nucleated HBB grows by recruiting free HBB proteins until it reaches a threshold level, at which point the HBB is considered complete [28, 29]. Complete HBBs do not recruit free HBB proteins but secrete FlgM, FliD, and FliC, which results in free *σ*^28^ and FliT within the bacterium. Depending on the HBB protein production rate and recruitment rate of incomplete HBBs, the bacteria are capable of growing multiple HBBs simultaneously. Let *h* be the concentration of free HBB proteins in the bacterium, and *b*_*i*_ be the number of HBB proteins in the *i*^th^ incomplete HBB. We then have

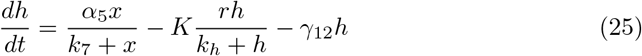

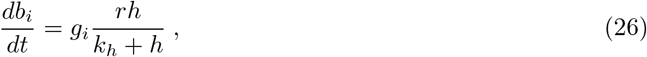

where *α*_5_ is the production rate of free HBB proteins, *k*_7_ is the Michaelis constant for HBB protein expression, *r* is the recruitment rate of free HBB proteins to incomplete HBBs, *k*_*h*_ is the Michaelis constant for HBB protein recruitment, *γ*_12_ is the degradation rate of free HBB proteins, and *g*_*i*_ is zero or one, depending on whether the *i*^th^ HBB is growing. *K* is the number of incomplete HBBs, 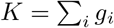. The nucleation and completion thresholds are *h*_*nuc*_ and *h*_*max*_, respectively.

#### Simulation methods

We use the parameters in Table 2 to simulate the model in MATLAB (2019a; MathWorks, Natick, MA).

**Table 2.** Model parameter descriptions and values.

| Parameter | Description | Value |
| --- | --- | --- |
| $\rho_1$ | Maximum production rate of FlhD<sub>4</sub>C<sub>2</sub> | 2.6 |
| $\rho_2$ | Production rate of RflP | 1.25 |
| $\alpha_1$ | Production rate of FliZ | 2 |
| $\alpha_2$ | Production rate of RflM | 1 |
| $\alpha_3$ | Production rate of FliT:FliD complex | 1 |
| $\alpha_4$ | Production rate of $\sigma^{28}$:FlgM complex | 1 |
| $\alpha_5$ | Production rate of *H* proteins | 1 |
| $\beta_1$ | Class III production of FliZ | 2 |
| $\beta_2$ | Production rate of FliC | 1 |
| $k_1$ | Michaelis constant for FliZ expression | 5 |
| $k_2$ | Michaelis constant for RflM expression | 1 |
| $k_3$ | Michaelis constant for FliT:FliD expression | 0.1 |
| $k_4$ | Michaelis constant for $\sigma^{28}$:FlgM expression | 0.1 |
| $k_5$ | Michaelis constant for Class III FliZ expression | 5 |
| $k_6$ | Michaelis constant for FliC production | 1 |
| $k_7$ | Michaelis constant for *H* protein expression | 5 |
| $\eta_1$ | Repression coefficient of RflM on FlhD<sub>4</sub>C<sub>2</sub> expression | 0.01 |
| $\eta_2$ | Repression coefficient of FliZ on RflP expression | 0.5 |
| $\kappa_1$ | Rate of binding of RflP and FlhD<sub>4</sub>C<sub>2</sub> | 1 |
| $\kappa_2$ | Rate of binding of FliT and FlhD<sub>4</sub>C<sub>2</sub> | 2 |
| $\gamma_1$ | Degradation rate of FlhD<sub>4</sub>C<sub>2</sub> | 0.1 |
| $\gamma_2$ | Degradation rate of FliZ | 0.1 |
| $\gamma_3$ | Degradation rate of RflP | 0.1 |
| $\gamma_4$ | Degradation rate of RflP:FlhD<sub>4</sub>C<sub>2</sub> complex | 0.01 |
| $\gamma_5$ | Degradation rate of RflM | 0.1 |
| $\gamma_6$ | Degradation rate of FliT | 0.1 |
| $\gamma_7$ | Degradation rate of FliT:FlhD<sub>4</sub>C<sub>2</sub> complex | 0.01 |
| $\gamma_8$ | Degradation rate of FliT:FliD complex | 0.01 |
| $\gamma_9$ | Degradation rate of $\sigma^{28}$:FlgM complex | 0.01 |
| $\gamma_{10}$ | Degradation rate of $\sigma^{28}$ | 0.1 |
| $\gamma_{11}$ | Degradation rate of FliC | 0.1 |
| $\gamma_{12}$ | Degradation rate of free *H* proteins | 0.01 |
| $\delta_1$ | ClpXP degradation rate of FlhD<sub>4</sub>C<sub>2</sub> in RflP complex | 0.1 |
| $\delta_2$ | ClpXP degradation rate of FlhD<sub>4</sub>C<sub>2</sub> in FliT complex | 0.1 |
| $r$ | Recruitment rate of free *H* proteins to incomplete HBBs | 0.1 |
| $k_h$ | Michaelis constant for recruitment of free *H* proteins | 1 |
| $\sigma$ | Total secretion rate through the HBBs | 0.06 |
| $k_s$ | Michaelis constant for total secretion through the HBBs | 1 |
| $h_{nuc}$ | HBB nucleation threshold of free *H* proteins | 15 |
| $h_{max}$ | HBB completion threshold of *H* proteins | 100 |

Parameter values were chosen to capture the desired dynamics of the system. Concentration and time both have arbitrary units.

We use all zero initial conditions, unless specified otherwise, and set *K* = 0 and *J* = 0. We then numerically integrate Eqs 17–26 until *h* reaches the nucleation threshold, *h*_*nuc*_, which indicates the first nucleation event. Using the state of the system just prior to nucleation and setting *h* → 0, *b*_1_ → *h*_*nuc*_, *g*_1_ → 1, *K* → 1, we numerically integrate the system again until *h* = *h*_*nuc*_ or *b*_*i*_ = *h*_*max*_. If *h* = *h*_*nuc*_, another HBB is nucleated, meaning *h* → 0, *b*_*j*_ → *h*_*nuc*_, *g*_*j*_ → 1, *K* → *K* + 1, where the *j*th HBB is newly nucleated. If *b*_*i*_ = *h*_*max*_, we complete construction of the *i*th HBB, meaning *b*_*i*_ → 0, *g*_*i*_ → 0, *K* → *K* − 1, *J* → *J* + 1. The process continues until the system reaches steady-state.

## Results

The bistability observed in the sub-system model is maintained in the extended model. Depending on the initial conditions, HBBs are either produced or not. We first assume an initial condition of zero for all components of the gene network. Due to basal production of FlhD_4_C_2_, there is some expression of Class II genes (Fig 7A), but an insufficient concentration of FlhD_4_C_2_ is accumulated to initiate HBB production (Fig 7B).

**Fig 7.**
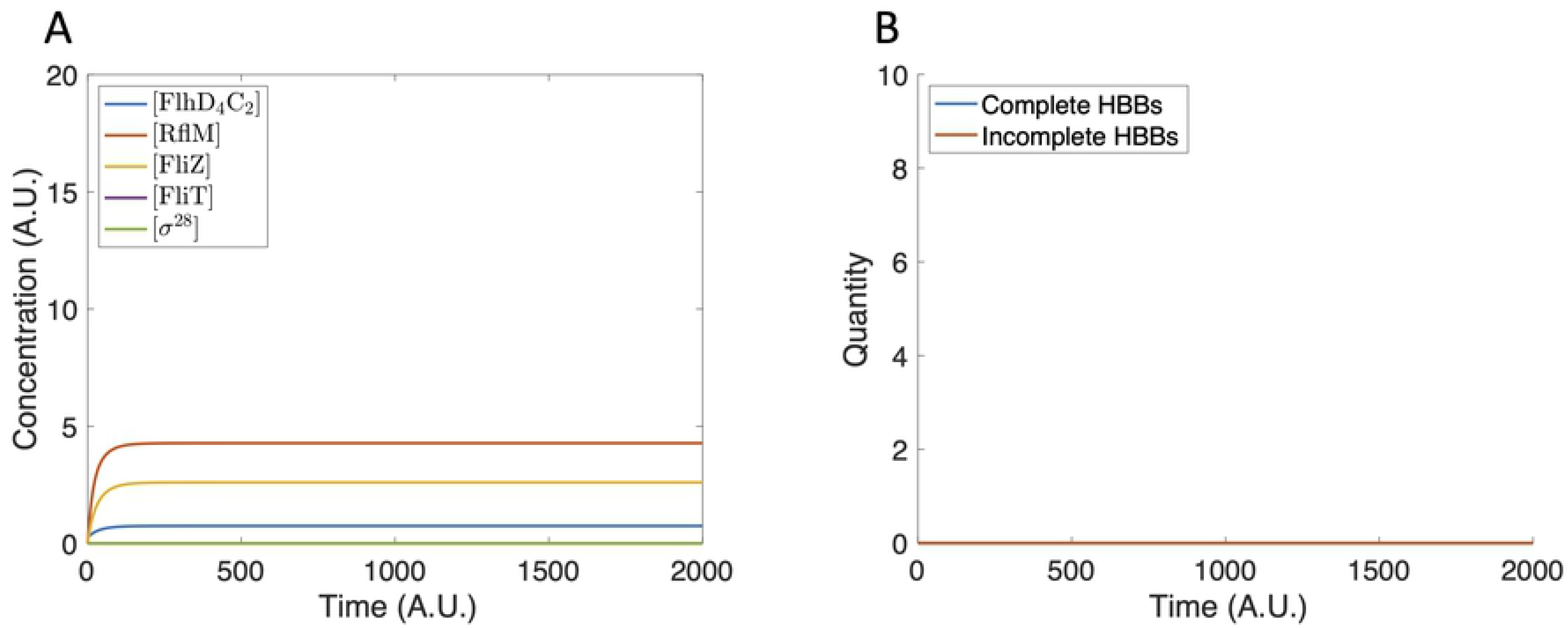
Model dynamics using zero initial conditions. (A) The protein concentration increases and reaches steady state. The protein concentrations are not large enough to initiate HBB production. (B) The tally of incomplete and complete HBBs remains at zero for all time.

In contrast, when we assume a small initial concentration of FliZ and FlhD_4_C_2_, FlhD_4_C_2_ accumulates sufficiently to initiate HBB production (Fig 8A), resulting in the production of seven HBBs (Fig 8B).

**Fig 8.**
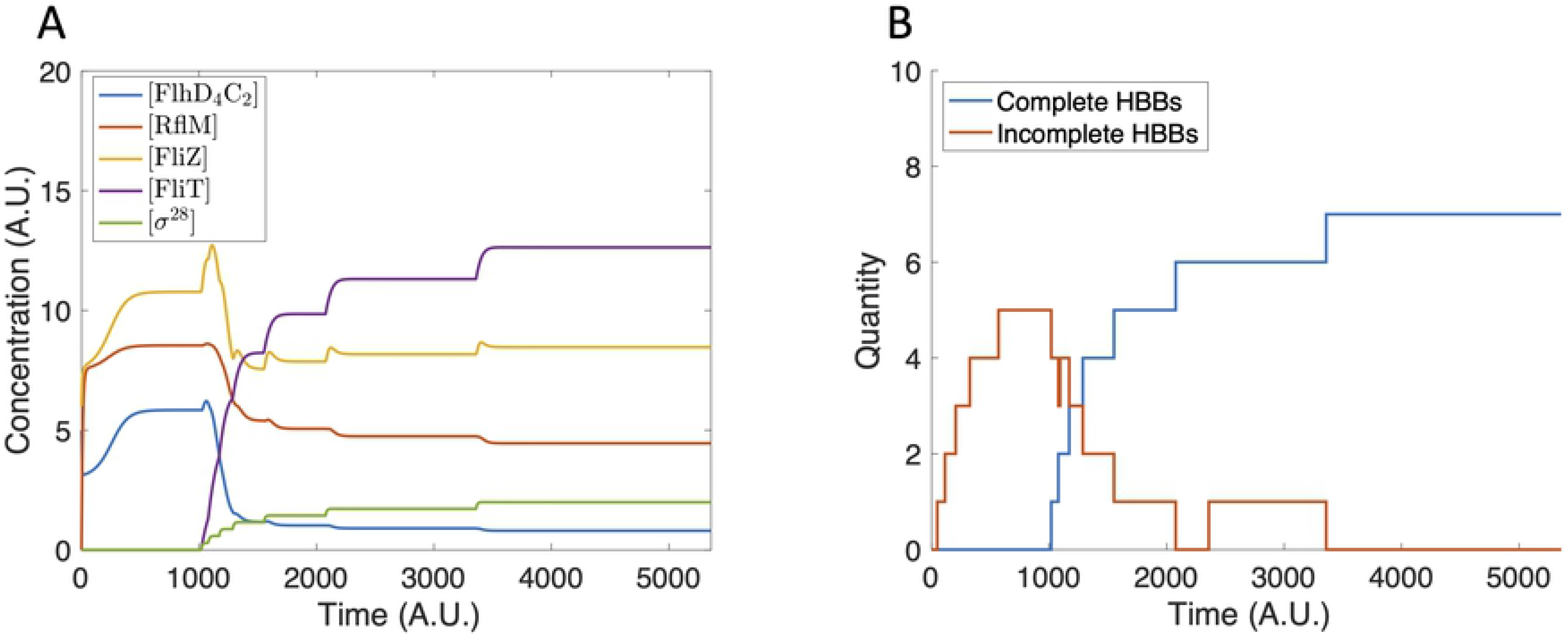
Model dynamics using [FlhD_4_C_2_](0)=5 and [FliZ](0)=6. (A) The protein concentrations increase until completion of the first HBB. The concentrations then decrease and reach a steady state that does not support further HBB nucleation. (B) Seven HBBs are produced.

The differing outcomes confirm that the FlhD_4_C_2_-FliZ-RflP feedback loop, as predicted by the sub-system model, determines if HBBs are produced. We then compute the separatrix in [FlhD_4_C_2_]-[FliZ] initial condition space to determine what initial conditions are necessary to stimulate HBB production (solid curve in Fig 9).

**Fig 9.**
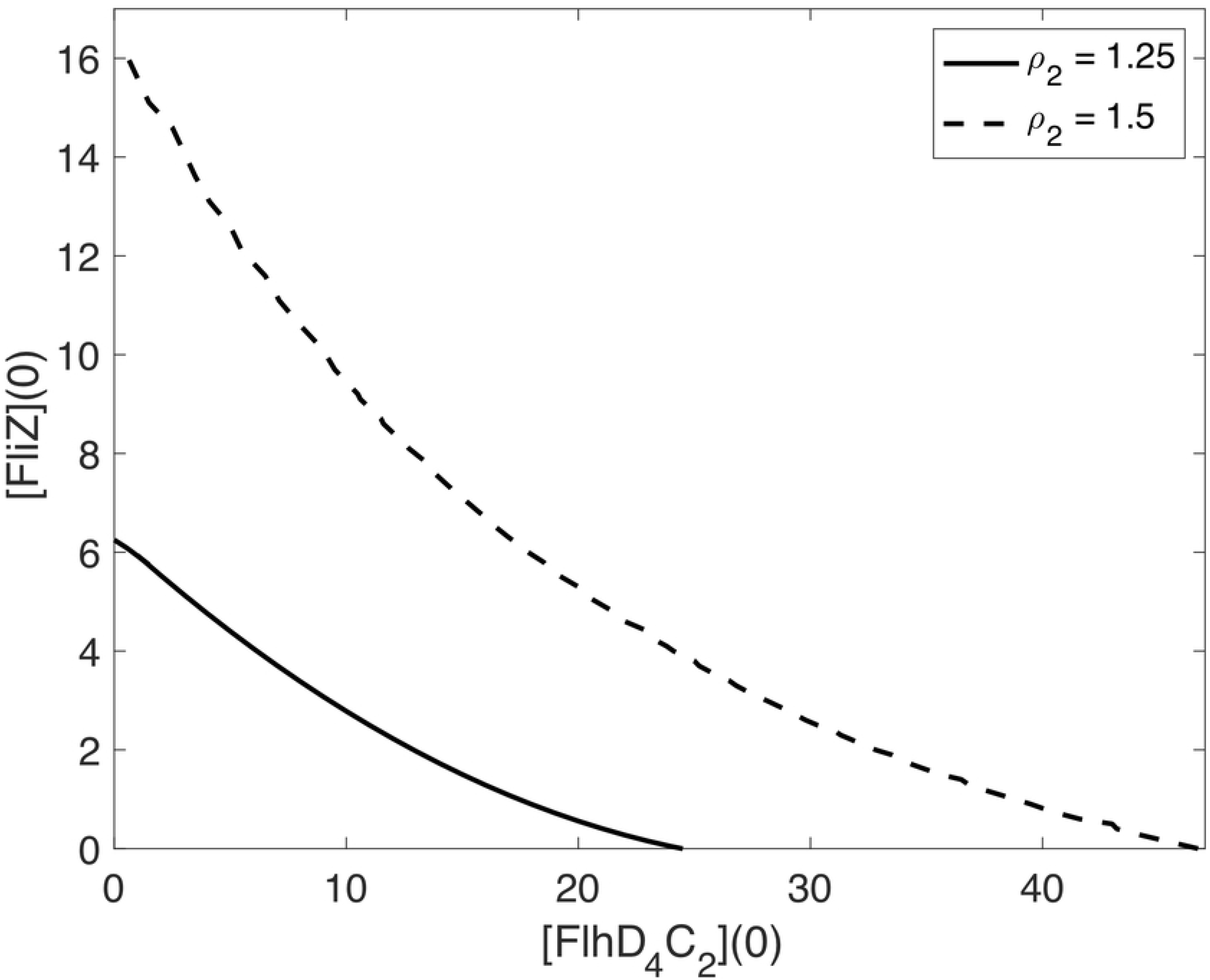
Separatrices in [FlhD_4_C_2_]-[FliZ] initial condition space. The solid curve indicates the *ρ*_2_ = 1.25 case and the dashed curve indicates the *ρ*_2_ = 1.5 case. Initial conditions to the right of the curve lead to HBB production, while initial conditions to the left of the curve do not.

We next consider the impact of nutrient availability on HBB production by changing the RflP production rate, *ρ*_2_. We use *ρ*_2_ as a proxy for environmental nutrient conditions in which increased nutrient availability decreases *ρ*_2_ and decreased nutrient availability increases *ρ*_2_. A 20% decrease in *ρ*_2_ results in the production of eight HBBs (Fig 10A), which is one more in comparison to the baseline case, whereas increasing *ρ*_2_ by 20% results in no HBB production (Fig 10B).

**Fig 10.**
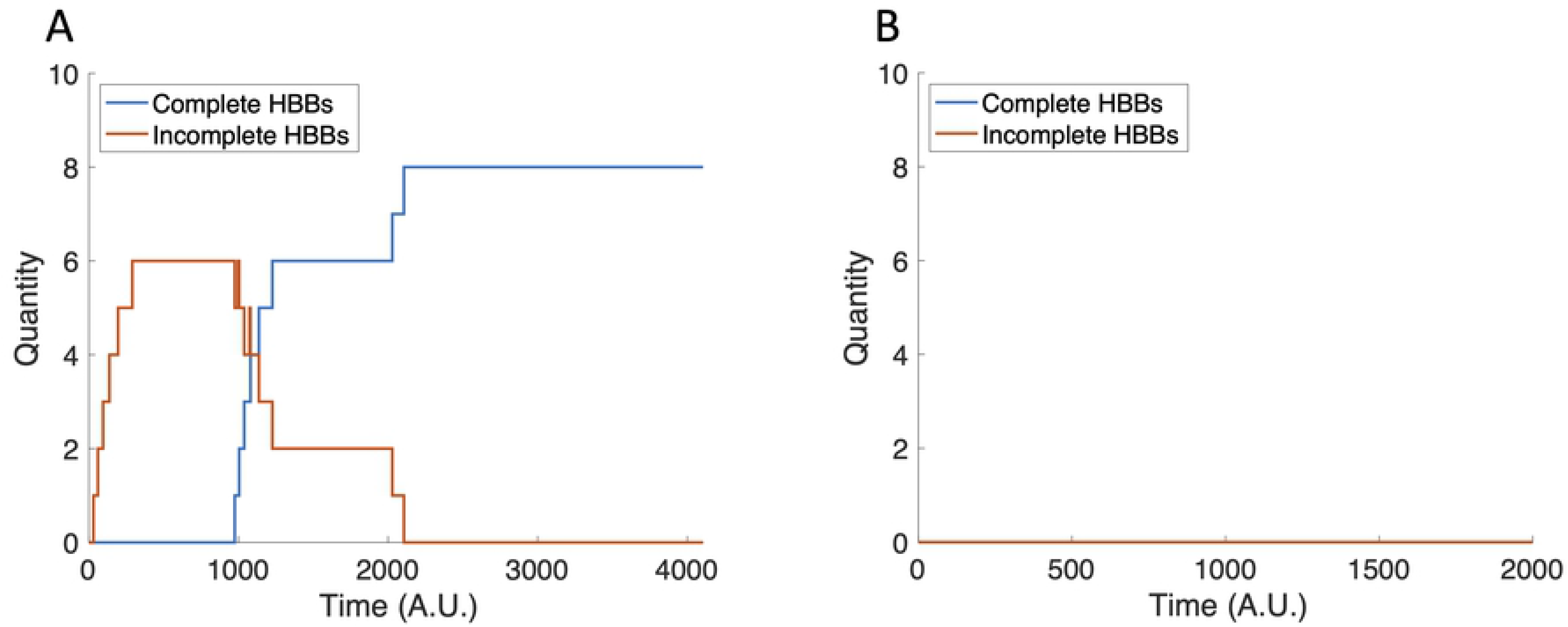
The tally of incomplete and complete HBBs with perturbed *ρ*_2_ values. (A) Eight HBBs are completed when *ρ*_2_ = 1. (B) HBBs are not produced when *ρ*_2_ = 1.5.

Since changing *ρ*_2_ results in a different number of HBBs produced, we know that *ρ*_2_ impacts the separatrix between no HBB production and HBB production. Increasing *ρ*_2_ by 20% shifts the curve to require larger initial concentrations of FlhD_4_C_2_ and FliZ to produce HBBs (dashed curve in Fig 9). A 20% decrease in *ρ*_2_ shifts the curve such that HBBs are produced under any biologically reasonable initial conditions (not shown).

We next investigate the impact of the FliT:FliD production rate, *α*_3_, on HBB production. Decreasing *α*_3_ results in more HBBs (Fig 11A), while increasing *α*_3_ results in fewer HBBs (Fig 11B), in comparison to the case with baseline *α*_3_.

**Fig 11.**
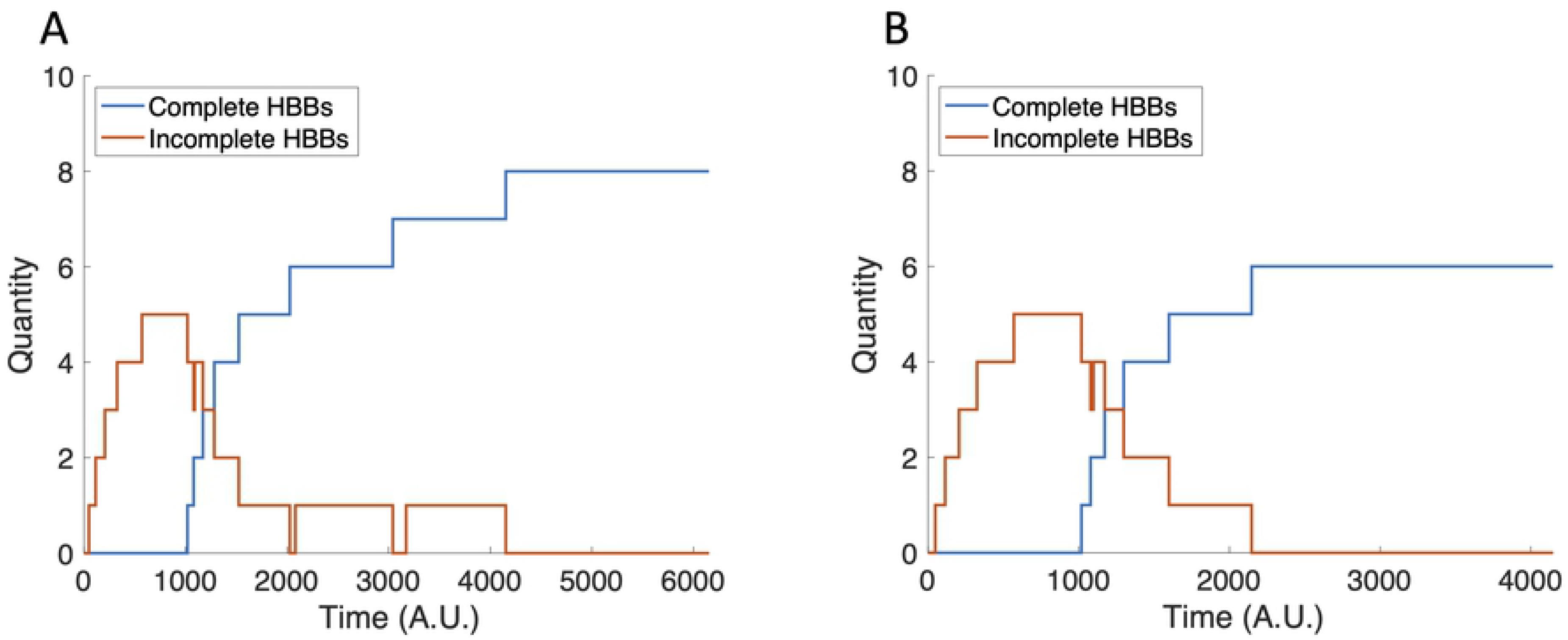
The tally of incomplete and complete HBBs with perturbed *α*_3_ values. (A) Eight HBBs are completed when *α*_3_ = 0.8. (B) Six HBBs are completed when *α*_3_ = 1.2.

The change in HBB production in response to changes in *α*_3_ suggests that the FliT:FliD concentration, and, consequently, the free FliT concentration, is a key factor in controlling the termination of HBB production. This is exemplified by setting *α*_3_ to zero (Fig 12). In this case, HBB construction continues indefinitely, which means the repressive activity of FliT on FlhD_4_C_2_ is required to stop HBB production.

**Fig 12.**
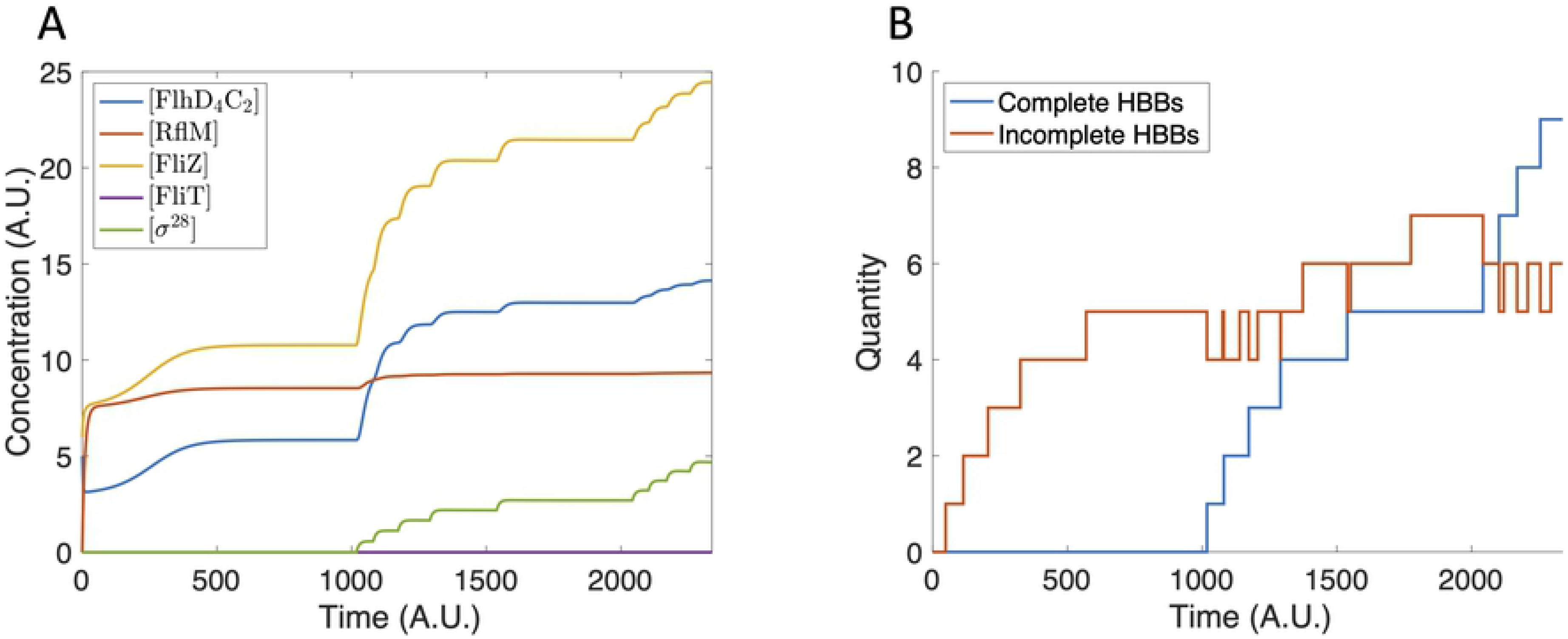
Model dynamics with *α*_3_ = 0. (A) The protein concentrations increase indefinitely and FliT concentration remains zero. (B) The tally of incomplete and complete HBBs increases indefinitely.

To further consider the impact of FliT on FlhD_4_C_2_ concentration, we recall the sub-system model (Eqs 1–5). Even though FliT is not explicitly included in the model, increasing the basal rate of degradation of FlhD_4_C_2_, *γ*_1_, can be interpreted as a proxy for the repressive activity of FliT on FlhD_4_C_2_. As described earlier, Fig 5A shows that FlhD_4_C_2_ concentration is bistable as a function of *γ*_1_. This means that as HBBs are completed and free FliT levels increase, *γ*_1_ increases, which results in a saddle-node bifurcation and collapse of the system to a single, low steady state value. Similarly, Fig 5B shows that the bistability of FlhD_4_C_2_ concentration as a function of *ρ*_2_ is sensitive to *γ*_1_. As *γ*_1_ increases, the system moves outside the region in parameter space that supports bistability. These results suggest that there is a threshold level of free FliT required to shut-down HBB production.

Finally, we investigate the role of Class III regulation of FliZ. In particular, we set the Class III production rate of FliZ, *β*_1_, to zero. Without Class III regulation of FliZ, the FliZ concentration drops dramatically following the first HBB completion (Fig 13A), and, as a result, only five HBBs are produced (Fig 13B). The reduction in the number of HBBs produced, in comparison to the baseline case, is due to the lack of HBB nucleation following the first HBB completion event. Following HBB completion, FliT activity decreases the concentration of FlhD_4_C_2_. This slows the production of FliZ, and, without Class III regulation, the FliZ concentration drops dramatically. The drop in FliZ concentration further contributes to the decrease in FlhD_4_C_2_ concentration. FlhD_4_C_2_ then drops to a level that does not support further HBB nucleation.

**Fig 13.**
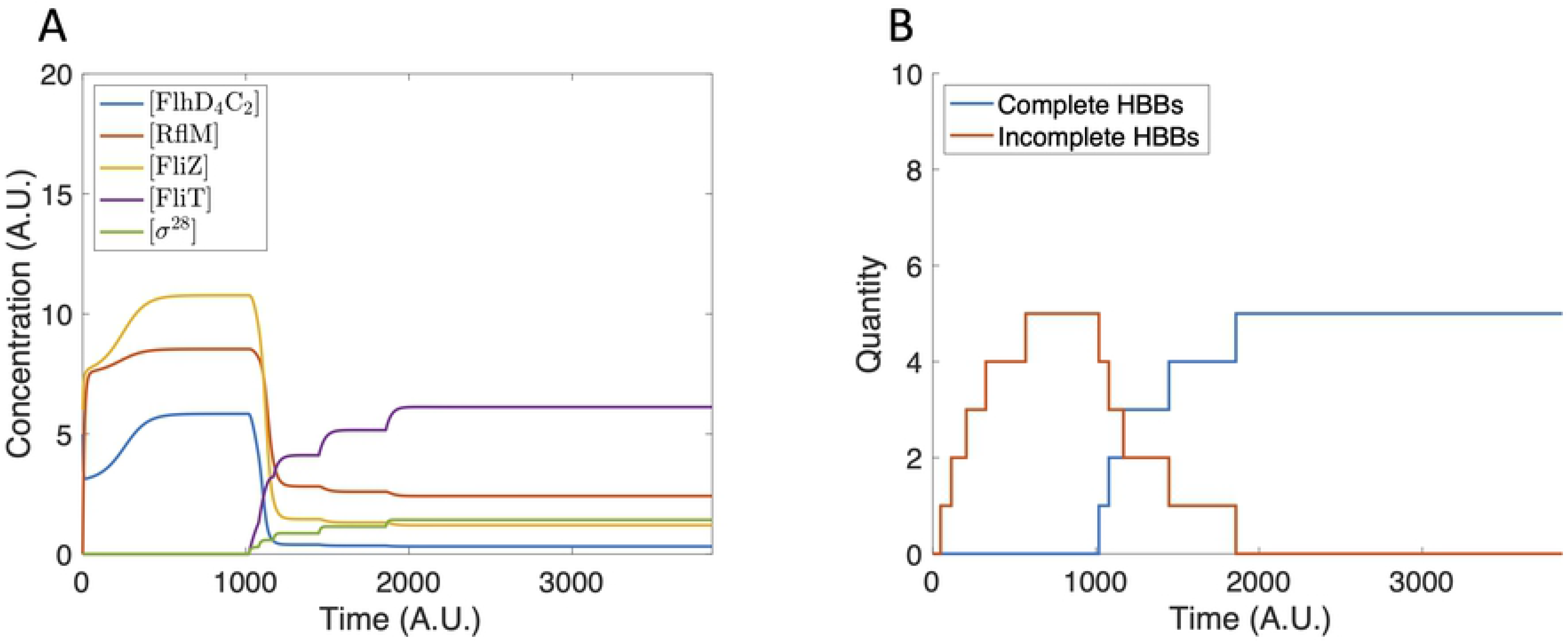
Model dynamics with *β*_1_ = 0. (A) The protein concentrations increase until the first HBB is completed. The concentrations then rapidly decrease to a level that does not support HBB nucleation. (B) Five HBBs are constructed. No nucleation events occur after completion of the first HBB.

Together, these results suggest that the FlhD_4_C_2_-FliZ-RflP feedback loop determines whether flagella production will be initiated, and the balance of FliT:FliD dynamics and Class III regulation of FliZ control the total number of flagella produced.

## Discussion

*S. enterica* is a common foodborne pathogen that impacts millions of people each year, and the flagellum is an essential component of *S. enterica* pathogenesis. Motility is critical in enabling bacteria to reach the site of infection and establish disease. Therefore, understanding how *S. enterica* regulate and construct flagella is of interest for disease prevention and treatment. An interesting feature of flagella regulation in *S. enterica* is the presence of a heterogeneous quantity of flagella per bacterium in a clonal population [3].

Here, we developed mathematical models of the gene network that regulates flagella construction to improve our understanding of flagella heterogeneity. Analysis of a sub-system model (Eqs 1–5) of the network identified the possibility of bistable FlhD_4_C_2_ concentrations. The bistability was determined by the production rates of FlhD_4_C_2_ and RflP, and could explain why some bacteria within a clonal population grow flagella while others do not.

We then extended the sub-system model to include the construction of HBBs and additional regulatory mechanisms. Simulations of this model (Eqs 17–26) suggest that FliT activity and Class III regulation of FliZ provide a counting mechanism by which the bacteria can control how many flagella are produced. Class III production of FliZ prevents underproduction of flagella, and FliT prevents overproduction. Hence, FlhD_4_C_2_ is regulated by multiple negative feedback loops to ensure that protein levels in the bacteria can be tightly controlled, and both under- and overproduction of flagella is avoided.

The *flhDC* operon is primarily transcribed from two promoters, P1_*flhDC*_ and P5_*flhDC*_, and many additional factors are shown to positively and negatively regulate *flhDC* expression, including HilD, RcsDBC, RtsB, and LrhA [21, 30–32]. Our model includes the main components involved in the expression of *flhDC* from P1_flhDC_. Expression from the P5_flhDC_, which is regulated via a FliZ-HilD positive feedback loop [21], plays a role in virulence rather than motility. Therefore, we focus on expression from P1_flhDC_. Future models could include expression from P5_flhDC_ and additional regulatory mechanisms.

The limited availability of data meant that parameter values were selected manually, rather than measured or estimated. This study investigated the possible behaviors of the gene network, and, thus, does not rely on accurate parameter estimates. As additional data becomes available, the parameter values could be updated. Further, due to a limited understanding of HBB construction, we have proposed a simple model of the process. As we learn more about HBB construction, the model could also be updated.

Our results explain the bimodality of flagella count data and how the bacteria regulate the total number of flagella produced. However, since the model is deterministic, it cannot capture the spread of 6–10 flagella in the count data. Future models could incorporate sources of stochasticity, such as low protein copy numbers, to understand the variability in the number of flagella. Further, the impact of cell division on population-level flagella distributions could also be considered.

This paper has proposed two mathematical models of the flagellar gene network in *S. enterica*. We predict that the FlhD_4_C_2_-FliZ-RflP feedback loop determines whether flagella are produced, and the combination of FliT activity and Class III regulation of FliZ determines the total number of flagella produced.

